# A dopamine-dependent decrease in dorsomedial striatum direct pathway neuronal activity is required for learned motor coordination

**DOI:** 10.1101/2021.06.07.447452

**Authors:** Stefano Cataldi, Clay Lacefield, N Shashaank, Gautam Kumar, David Sulzer

## Abstract

It has been suggested that the dorsomedial striatum (DMS) facilitates the early stages of motor learning for goal-directed actions, whereas at later stages, control is transferred to the dorsolateral striatum (DLS), which enables the motor actions to become a skill or habit. It is unknown whether these striatal regions are simultaneously active while expertise is acquired during skill learning. To address this question, we developed the “treadmill training task” to track changes in mouse locomotor coordination during practice at running that simultaneously provides a means to measure local neuronal activity using photometry. We analyzed body position and paw movement to evaluate changes in motor coordination over practice sessions on the treadmill using DeepLabCut and custom-built code. By correlating improvements in motor coordination during training with simultaneous neuronal calcium activity in the striatum, we found that DMS direct pathway neurons exhibited decreased activity as the mouse gains proficiency at running. In contrast, direct pathway activity in the DLS was similar throughout training and did not correlate with learned skill proficiency. Pharmacological blockade of D1 dopamine receptors in these subregions during task performance confirmed that dopamine neurotransmission in the DMS direct pathway activity is necessary for efficient motor coordination learning, while dopamine signalling in the DLS is important for both coordination learning and maintenance of the acquired skill. These results provide new tools to measure changes in fine motor skills during simultaneous recordings of brain activity, revealing fundamental features of the neuronal substrates of motor learning.

## Introduction

Neural circuits in the striatum are central to basal ganglia functions, including the learning and organization of motor skills. The development of techniques to record the activity of specific populations of neurons in the striatum along with pharmacological or genetic models of motor disorders has revealed new details about the pathways that underlie motor performance and motor learning. The dorsal portion of the striatum has been found to be particularly important for expressing automatic actions (Yin et al., 2009). In rodents, this region is typically separated into two major regions: the dorsomedial striatum (DMS) and the dorsolateral striatum (DLS), corresponding to the human caudate and putamen, respectively. The DMS receives afferents from prefrontal and associative cortices, while the DLS mostly receives input from sensorimotor cortical areas (Cataldi et al., 2021; Graybiel, 2008; Hunnicutt et al., 2016).

*In vivo* electrophysiological recordings indicate that the DMS is engaged in early training on a rotarod (Lingawi & Balleine, 2012; Yin et al., 2009), a task commonly used to study motor coordination, while the DLS is particularly active later in training. A recent study that analyzed corticostriatal synaptic responses in brain slices prepared from mice trained on the rotarod indicated a transient decreased postsynaptic response by DMS striatal spiny projection neurons (SPNs) following training (Badreddine et al., 2022). However, single unit recording during a lever press task showed a similar activity pattern in DLS and DMS after this skill was acquired (Vandaele et al., 2019).

SPNs constitute the majority of striatal neurons (Silberberg & Bolam, 2015) and are generally classified as either D1 dopamine receptor expressing SPNs that form the direct pathway (D1-SPNs) or D2 dopamine receptor expressing SPNs that form the indirect pathway (D2-SPNs; Gangarossa et al., 2013; Gerfen et al., 1990; Kawaguchi, Wilson, & Emson, 1990; Wall et al., 2013). D1-SPNs axons project to basal ganglia output nuclei, particularly the globus pallidus internal segment and substantia nigra pars reticulata, while D2-SPNs project to the globus pallidus external segment, thus making indirect connection with output nuclei (Albin, Young, & Penney, 1989; DeLong, 1990). According to these classical models, activation of the direct pathway releases neurons in the motor thalamus from inhibition, thus promoting movement. Consistent with these models, optogenetic activation of the D1-SPNs increases locomotion and reduces freezing (Kravitz et al., 2010).

The activity of D1-SPNs has been studied at the level of the initiation, termination, and velocity of a specific motor action (Jin, Tecuapetla, & Costa, 2014), but there has been little investigation of the role of such circuits during motor learning, an important and dynamic phase of behavioral performance (Cataldi et al., 2021). Computational analysis of behavioral and physiological data during task learning promises to provide a means to elucidate the basis of learning complex motor actions, such as running, under a range of experimental conditions during this crucial phase.

Here we introduce a paradigm to measure fine changes in motor coordination *in vivo* as a mouse acquires the ability to run on a motorized treadmill while simultaneously recording activity of direct pathway neurons in the DLS or DMS, in order to study the neural and behavioral changes that occur across early and late stages of learning. We combine a well-established pose tracking method, DeepLabCut (Mathis et al., 2018), with customized Python-based software to analyze and interpret the behavioral data at different phases of learning. We then correlate the results with changes in calcium activity measured by fiber photometry within striatal regions over training. Our results indicate that DMS activity is reduced over training in running on the treadmill, while DLS activity is similar during early and late stages of learning. Furthermore, pharmacological inhibition of D1-SPNs by D1 antagonists in the DLS affects performance in both early and late training, while similar inhibition of neurons in the DMS delays skill acquisition without affecting performance after the task is learned.

## Materials and Methods

### Animals

Wild-type and transgenic mice expressing Cre-recombinase in D1-expressing SPNs (D1-cre) were used for these experiments. All animal procedures were approved by the Institutional Animal Care and Use Committee of the New York State Psychiatric Institute. Experiments were performed on 3 to 6-month-old male and female mice. C57BL/6 mice from Jackson Laboratory (Jax #000664) were used as control animals. D1-Cre transgenic mice were obtained from MMRC/Jax (ey262 D1- cre tg-drdla-cre) and crossed with C57BL/6 mice.

### Viral expression of GCaMP6f and fiber implants

To achieve neuronal subtype-specific expression of the genetically encoded calcium sensor GCaMP6f in direct pathway SPNs, AAV vectors containing Cre-inducible GCaMP6f (AAV.9.Syn.Flex.GCaMP6f.WPRE.SV40, Addgene; TIter≥2.1×13 GC/ml) were injected into the right DLS or DMS by stereotaxic surgery. During the surgery, a small skull craniotomy (1 mm × 1 mm) above the injection site was opened with a dental drill. A glass pipette attached to a Nanoject II (Drummond Scientific) was filled with the GCaMP6f AAV and lowered to target locations in the dorsomedial or dorsolateral striatum (tip coordinates from bregma: AP −0.5 mm, ML +2.5 DV −3.2 mm for DLS, AP +1.2 mm ML −0.8 mm DV −2.8 mm for DMS). A total volume of 150 nl AAV vector per site was injected over 10 minutes. The pipette was left in place for 5 more minutes before removal. A 4 mm long 300 µm diameter optic fiber (Doric Lenses; MFC 300/370-0.22_4 mm_MF2.5_FLT) was subsequently lowered to the same coordinates. The skull was then covered with dental acrylic to secure the optic fiber in place. Alternatively, opto-fluid cannulas (Doric Lenses; OmFC_MF1.25_300/370-0.22_4.2_FLT_4) were implanted for simultaneous GCaMP6f recording and local microinjection of drug or saline. Animals were allowed to recover and photometry experiments were performed four weeks after surgery, for optimal viral expression. Three to five days before running on the treadmill, animals were placed in an open field chamber to record baseline calcium signals and habituate to the tethered optical fiber.

### Custom-built treadmill

To precisely control the animal’s locomotion, mice were placed on a custom motorized treadmill (Supplementary Figure 1A). The treadmill consisted of a 1 m clear belt stretched between two 3 inch diameter acrylic wheels on an aluminum frame (8020.net). Treadmill speed was controlled by an Arduino-based system (OpenMaze.org, OM4 board) that adjusted the speed of a 12 V gear motor attached to the axle of one of the treadmill wheels through pulse-width modulation (PWM). Belt speed was measured using a quadrature rotary encoder (Digikey #ENS1J) attached to the other axle and decoded by the Arduino. The Arduino/OpenMaze setup was also used to send synchronization pulses to coordinate behavior with video recording and fiber photometry. Mouse movement on the treadmill was constrained by placing the mouse into a 6 inch-long clear acrylic box over the center of the treadmill and covering the entire width of the belt, which ensured the animal was walking along the belt to avoid being forced into the back wall during belt movement. An acrylic mirror was fixed at a 45° angle under the clear acrylic box and mesh belt to allow high speed videography of the animal’s locomotion from lateral and ventral viewpoints simultaneously.

### Treadmill Training Task

Mice undergoing experimental procedures were weighed prior to testing. All tests were performed in the morning between 9:30 am and 1:30 pm, during the animals’ typical awake phase. All animals were assessed at 4-6 months of age, and all experimentation and analysis were conducted with the experimenter blinded to the location of the implant or condition. After 3-day familiarization to experimenter handling, mice underwent the following training paradigm.

Mice were placed on the treadmill within the clear inbuilt open box and were left free to explore the environment for 30 seconds. After the 30 second baseline, the treadmill movement was activated through the Arduino system at a low speed of 3 m/min, with an increment every 60 seconds up to 12 m/min (total of 5 speeds; intermediate speeds are 6 m/min, 8 m/min, and 10 m/min; Figure 1A). After 5 minutes of running, the treadmill was turned off for a final baseline recording of 30 seconds. Finally, animals were removed from the treadmill and placed back into their home cage. This process was then repeated for 12 consecutive days, including home cage baseline recording and pre/post-running treadmill recording epochs. Animal weight was recorded every day after training and showed no significant changes (Supplementary Figure 1B). The Arduino source code is freely available at github.com/DSulzerLab/treadmill.

**Figure 1.**
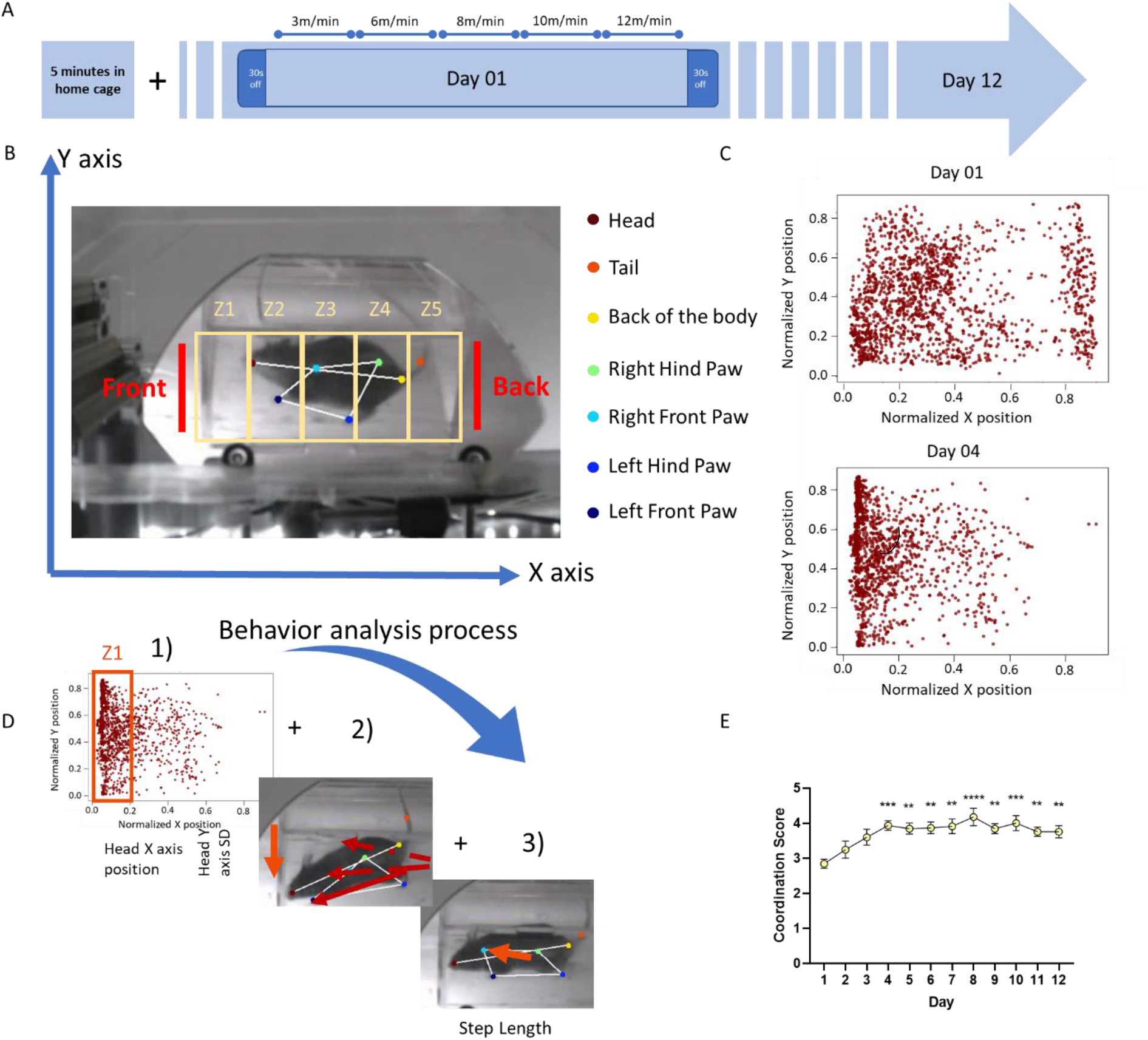
A) Timeline of behavioral protocol in the Treadmill Training Task. On each training day, a mouse is connected to the photometry patch cord in its home cage and a 5-minute calcium baseline signal recorded. After this habituation period, the mouse is placed on the treadmill and allowed to explore the environment for 30 seconds. The treadmill motor is then started at a velocity of 3 m/min, with the speed increased every 60 seconds up to 12 m/min, after which the treadmill is turned off for another 30 seconds. This protocol is repeated for 12 consecutive days. B) Schematic of the behavioral analysis. Positions of mouse body parts are obtained by analysis with DeepLabCut. For head position, the ventral 2D field of view are divided into five zones and the probability of the head of the mouse to be in each zone calculated. Color code for the body part is indicated in the legend on the right. C) Samples of head positions on day 1 (top) and day 4 (bottom) for a sample control mouse. Each point represents the head position in one frame and all frames from one video/session are overlapped. On day 1, the head position was frequently towards the rear of the field, indicating the mouse was falling behind the belt speed and often hitting the back wall. By day 4, the animal was able to keep up with the moving belt, and so the position of the head was more consistently towards the front. D) Process for obtaining the mouse motor coordination score. Motor coordination score is calculated from the expected value of the head position in the 5 zones, position along the Y axis (calculated from standard deviation of Y axis value), and step length (distance between steps for each paw), as detailed in the Methods. E) Motor coordination score for a cohort of control animals. Control mice show significant improvement in the coordination score over the 12 days of testing (n=9; 1-way ANOVA \**p*<0.05, *Bonferroni’s multiple comparisons test* as detailed in main text).

### Photometry recording

Calcium signals were recorded using a commercial fiber photometry apparatus (Doric Lenses). The system consisted of a console connected to a computer, a four-channel programmable LED driver, and two LEDs at 405 and 465 nm connected to fluorescence dichroic mini cubes and photometric multipliers tubes (PMTs). The 405 nm wavelength is the isosbestic point for GCaMP, a point where fluorescence does not change depending on calcium concentration (Supplementary Figure 2A). Detection at this wavelength was used to remove background noise (movement artifact or GCaMP auto-fluorescence). The 465 nm excitation provides detection of GCaMP signal where the intensity of fluorescence is proportional to cytosolic calcium concentration.

Calcium signals from all animals were recorded for 5 minutes as the mice explored their home cage prior to treadmill testing. This signal is used as a baseline to evaluate whether there is any loss of signal over the several days of testing, as well as to habituate the animal to the optical fiber.

### Calcium signal analysis

Calcium signals were processed using custom-built Python code to remove background noise and detect individual calcium peak events. The process consisted of 1) down sampling the signal to 30 samples/sec to match the sampling rate of behavioral recording, 2) normalization of the data and removal of background noise, 3) identification and quantification of peak events (events count and event amplitude). The source code is freely available at github.com/DSulzerLab/Calcium_peaks_analysis.

### Pharmacological experiments

SCH39166 was dissolved in 100% DMSO and then diluted to 1% DMSO concentration with saline (0.9% NaCl) containing 1 mM rhodamine (Rhodamine B Base; Aldrich 234141-10G), for a final concentration of 48.13µM SCH39166 (total injected 5.7 ng).

The mice were injected locally in the DMS or DLS with 300 nl SCH39166 (Tocris), a selective D1-antagonist, 1 mM at a rate of 100 nl/min using a microinjection pump (UltraMicroPump III, World Precision Instruments) through an injection cannula attached to the photometry fiber (Doric Lenses, Fl_OmFC_ZF_100/170_4.2) while running on the treadmill. Local injection was confirmed by recording the 565 nm signal from the rhodamine dye (Supplementary Figure 3). An additional cohort of control animals was injected with saline with rhodamine to ensure that the drug’s effect was not due to the injection or presence of rhodamine.

### Video Analysis

Behavioral experiments were recorded using a high speed USB webcam (Sony PS3eye) and commercial video acquisition software (Kinovea), synchronized to the Arduino/OpenMaze behavior system with an infrared LED, and analyzed *post-hoc* using DeepLabCut (Mathis et al., 2018) to track movement of individual body parts (the four paws, head, rear of the body, and tail were labeled; data was processed using Google Colaboratory; approximately 30,000-40,000 iterations were sufficient for good quality tracking).

The DeepLabCut results were subsequently processed with custom-built Python code to return a “motor coordination score” from 1 to 5, with 1 representing poor and 5 representing excellent coordination. The coordination score was computed using three values: the expected value of the mouse head’s X axis position, the standard deviation (SD) of the mouse head’s Y axis position, and the average step length of the four individual paws. First, the expected value of the mouse head’s X axis position was computed by dividing the box into five zones, with zone 1 being the frontmost and zone 5 the rearmost (Figure 1B), and calculating the probability of the head being positioned in a specific zone. Second, the SD of the mouse head’s Y axis position was computed to identify the sideways movement. Third, the step length was computed using the Python package “traja” (Shenk et al., 2021), which uses spatiotemporal animal tracking data to estimate parameters such as the length of each step, straightness (running along a straight line), and speed. The average step length for each paw was then computed, and the “paw stability” determined by calculating the step length’s distance from the median of the average step length measurements. Along with the paw stability, the percentiles of the expected values and the Y axis SD were computed, and these three values were given weights of 20/40/40, respectively. The percentiles of the weighted average were scaled between 1 to 5 to determine an overall coordination score. We arbitrarily chose the head position for this analysis, although position of either paw can also be used. Parallel analysis of the likelihood of paw placement over training demonstrates a higher fraction of steps within specific coordinates that are closer to the front of the box (Supplementary Figure 4).

Manual scoring of the videos under blinded conditions confirmed the validity and reproducibility of the test and analysis method. The source code is freely available at github.com/DSulzerLab/treadmill.

### Immunohistochemistry

After the completion of the behavioral experiments, mice were terminally anesthetized (euthasol 240 mg/kg, 200 µl i.p.) and intracardially perfused with PBS then 4% paraformaldehyde (PFA). Brains were extracted and post fixed overnight (4% PFA, 4°C). Coronal slices (100 µm) were obtained by vibratome (Leica VT 1200). Sections were rinsed with 0.6% Triton-X in 1x PBS (PBST; 6×60 min) and blocked in 10% normal donkey serum (NDS) in PBST (60 min, RT). Primary antibodies: chicken polyclonal GFP (ab13970 Abcam; 1:500) and rabbit polyclonal anti-tyrosine hydroxylase (TH; ab152 Abcam; 1:500), were applied in 2% NDS in PBST (48 h 4° C) prior to washing (6×60 min PBST) and secondary incubation with species specific Alexafluor IgG secondary antibodies (60 min RT, Invitrogen; 1:500). Tissues were washed again in 0.1% PBST (6 × 60 min), then mounted using DAPI Fluoromount-G® (0100-20, SouthernBiotech). Images were acquired using a 20x oil objective on an Olympus microscope (see sample images in Supplementary Figure 5).

### Statistics and data reporting

Data are presented throughout as mean ± SEM where n is the number of animals. Comparisons were conducted by 1- or 2-way ANOVA with appropriate *post-hoc* tests detailed in the text, using Prism 9.0 (GraphPad, San Diego California USA).

## Results

### Treadmill running

To explore motor coordination learning during locomotion, we designed a custom-made motorized treadmill that allows detection of fine improvement in paw mobility as a mouse learns to run on the treadmill at a range of velocities (referred as the Treadmill Training Task; Figure 1B and Supplementary Figure 1A).

A cohort of wild-type (WT) mice were initially examined. On the first day of the protocol, animals were placed on the treadmill and allowed to freely explore the environment for 30 seconds. After the 30 second baseline, the treadmill was activated using the Arduino system. After 5 minutes of running, the treadmill was turned off for a final baseline recording of 30 seconds (Figure 1A). No difference in behavior or calcium signals was found between the 30 second periods before and after the running time, and data from both off periods were merged and reported as “off-time.” This protocol was repeated for 12 consecutive days to track changes in the mouse running motor coordination.

The videos were analyzed with the machine-learning based videographic analysis platform DeepLabCut (DLC, Mathis et al., 2018), which allowed us to track paw movements in space and time. DLC provides X and Y coordinates of the 2D plane of the video for each body part of interest across individual frames (Figure 1B & C and Supplementary Figure 4). Results obtained by DLC were processed through custom-built R and Python scripts to compute a “motor coordination score” (see Methods) using variables such as the mouse head’s position within the box zones (X axis), the mouse’s sideways movement along the treadmill (Y axis SD), and the mouse paw coordination on the treadmill (average step length). In the initial days of training, mice had a significantly lower motor coordination score than during later sessions (Figure 1E), presumably as they had not been exposed to the treadmill before and displayed poor coordination during running. For example, the task naïve mice would tend to fall behind and make longer steps trying to catch up with the running tape. Moreover, naïve animals ran sideways rather than along a straight line (Supplementary Figure 4B). On later days of training, as mice accrued competency in running and coordination improved, their score improved (Figure 1E) and their likelihood of stepping in the same region increased (Supplementary Figure 4B). Control animals trained on the treadmill showed a significantly higher coordination score within the fourth day of training. By day 4, all mice performed significantly better than on the first training day (Figure 1E; 1-way ANOVA \**p*<0.05, *Bonferroni’s multiple comparisons test* day 4 *p*<0.001, day 5 *p*<0.01, day 6 *p*<0.01, day 7 *p*<0.01, day 8 *p*<0.001, day 9 *p*<0.01, day 10 *p*<0.001, day 11 *p*<0.01, day 12 *p*<0.01).

### DMS and DLS D1-SPNs calcium activity during running

We next explored the role of the direct pathway in skill acquisition by recording calcium signals from D1-SPNs in either the DMS or DLS. Recording of baseline D1-SPNs activity in the DMS show no significant changes over the 12 days (Figure 2B; 1-way ANOVA *p*=0.28). Similarly, no changes were found in D1-SPNs activity in the DLS (Figure 2B; 1-way ANOVA *p*=0.52).

**Figure 2.**
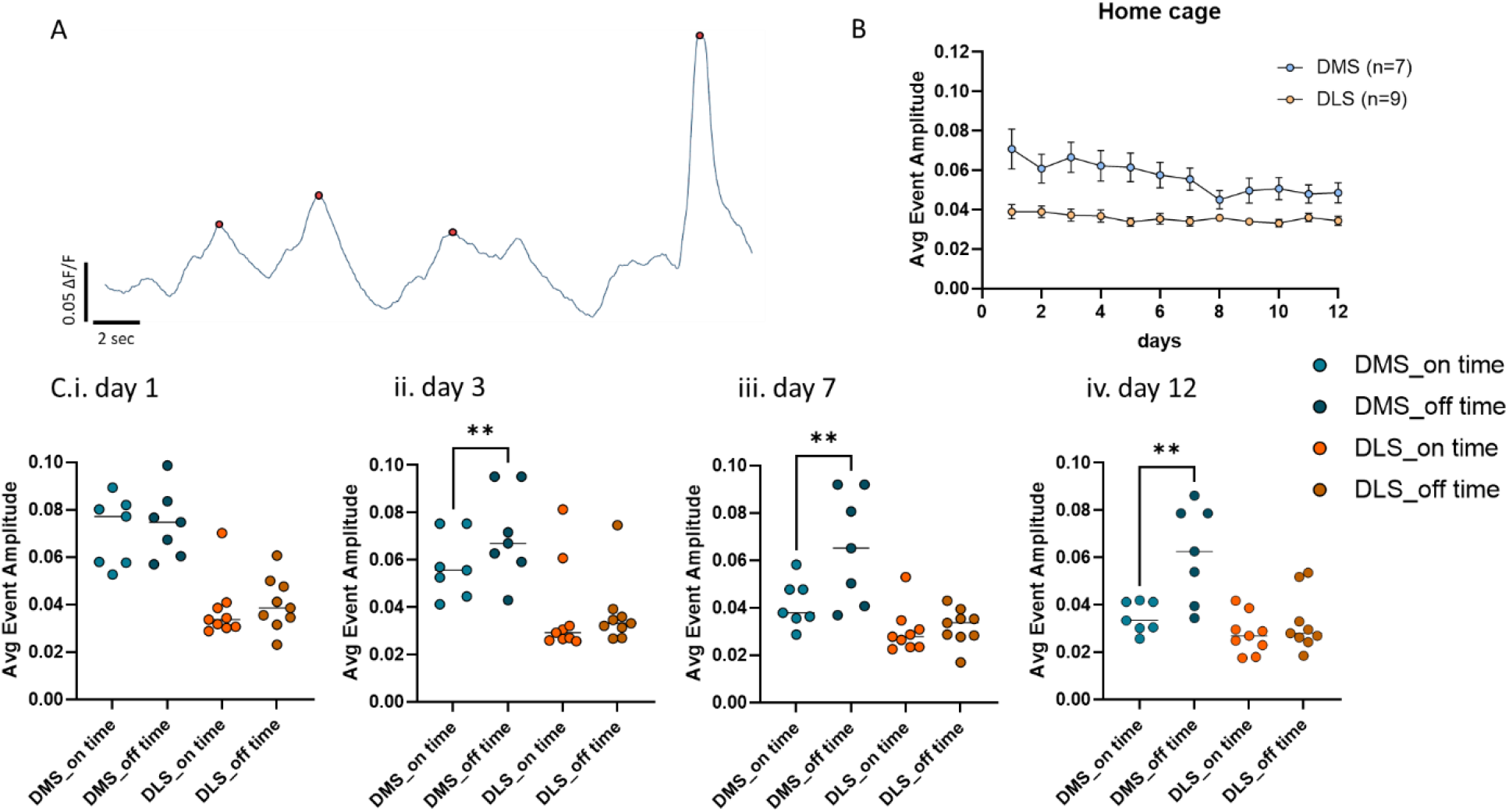
A) Sample trace of D1-SPNs calcium signals recorded in the DMS of a mouse running on the Treadmill Training Task. Detected calcium events are indicated by red dots. B) Average event amplitude of D1-SPNs in the DMS (light blue; n=7) and DLS (light orange; n=9) from the daily 5 minutes home cage baseline recording prior to treadmill testing. There is no significant decrease in event amplitude over the 12 days (1-way ANOVA *p*=0.28 for DMS and *p*=0.52 for DLS; n=6 and n=9 respectively). C) Average event amplitude from treadmill recording. Running epochs at all treadmill speeds are averaged together and referred to as “on time”. The 30 seconds prior to running and after the treadmill is turned off are averaged together and referred to as “off time”. Day 1 (C.i.), day 3 (C.ii.), day 7 (C.iii.), and day 12 (C.iv.) are shown. There are no changes between on and off time for D1-SPNs recorded from the DLS (n=6; day 01: 1-way ANOVA *p*=0.43, *Bonferroni’s multiple comparisons test p*=0.14 DLS on-time vs. DLS off-time; day 03: 1-way ANOVA *p*<0.01, *Bonferroni’s multiple comparisons test p*=0.06 DLS on-time vs DLS off-time; day07: 1-way ANOVA *p*<0.01, *Bonferroni’s multiple comparisons test p*=0.63 DLS on-time vs DLS off-time; day 12: 1-way ANOVA *p*<0.0001, *Bonferroni’s multiple comparisons test p*=0.95 DLS on-time vs DLS off-time). DMS D1-SPNs activity during on-time is comparable to off-time on day 1 (n=6; 1-way ANOVA *p*=0.43, *Bonferroni’s multiple comparisons test p*=0.79 DMS on-time vs DMS off-time) and becomes significantly lower than off-time activity starting on day 3 (1-way ANOVA *p*<0.01, *Bonferroni’s multiple comparisons test* \**p*<0.05 DMS on-time vs DMS off-time), and then on day 7 (1-way ANOVA *p*<0.01, *Bonferroni’s multiple comparisons test* \**p*<0.05 DMS on-time vs DMS off-time) and day 12 (1-way ANOVA *p*<0.0001, *Bonferroni’s multiple comparisons test* \*\**p*<0.01 DMS on-time vs DMS off-time), suggesting that activity in this region is lower once the skill is acquired.

During treadmill running, GCaMP6f-injected mice performed equally to control animals (Supplementary Figure 2B; 1-way ANOVA \**p*<0.05 for DMS mice and \*\**p*<0.01 for DLS mice), indicating that the surgery, viral expression, and implant did not affect the ability to run on the treadmill. We then correlated brain activity with the coordination score to examine neural changes as the mice gained proficiency at running on the motorized treadmill.

On day 1 of training, all mice showed similar average calcium event amplitude between the running (on-time; independently of the speed of the treadmill) and pre/post-running epochs in which the treadmill was off (off-time; Figure 2C.i. 1-way ANOVA *p*=0.43, *Bonferroni’s multiple comparisons test p*=0.14 DLS on-time vs DLS off-time; 1-way ANOVA *p*=0.43, *Bonferroni’s multiple comparisons test p*=0.79 DMS on-time vs DMS off-time). Similarly, event rates were comparable between baseline recording and while on the treadmill (on-time and off-time, Supplementary Figure 2C & D; 2-way ANOVA Time x Column Factor *p*=0.82 for DMS and *p*=0.77 for DLS).

Interestingly, DMS D1-SPNs calcium signals displayed a significant decrease in overall event amplitude beginning on day 3 while running (on-time) compared to off-time levels (Figure 2C.ii.; 1-way ANOVA *p*<0.01, *Bonferroni’s multiple comparisons test* \**p*<0.05 DMS on-time vs DMS off-time), which remained consistent throughout the 12 days of testing (Figure 2C.iii. & iv.; 1-way ANOVA *p*<0.01, *Bonferroni’s multiple comparisons test* \**p*<0.05 DMS on-time vs DMS off-time for day 7; 1-way ANOVA *p*<0.0001, *Bonferroni’s multiple comparisons test* \*\**p*<0.01 DMS on-time vs DMS off-time for day 12). Peak counts do not show the same trend (Supplementary Figure 2C; 2-way ANOVA Time x Column Factor *p*=0.82).

Conversely, D1-SPN calcium signals recorded from the DLS showed no significant difference in either calcium event amplitude or rate between running time and off-time over training, (Figure 2C; day 03: 1-way ANOVA *p*<0.01, *Bonferroni’s multiple comparisons test p*=0.06 DLS on-time vs DLS off-time; day 07: 1-way ANOVA *p*<0.01, *Bonferroni’s multiple comparisons test p*=0.63 DLS on-time vs DLS off-time; day 12: 1-way ANOVA *p*<0.0001, *Bonferroni’s multiple comparisons test p*=0.95 DLS on-time vs DLS off-time; Supplementary Figure 2D, 2-way ANOVA Time x Column Factor *p*=0.77 for DLS).

### Inhibition of D1-SPNs with a D1-receptor antagonist

We hypothesized that DMS changes over training are important for skill acquisition, while DLS activity is involved with performance. To investigate this, we locally injected SCH39166, a selective D1-antagonist, into the DMS or DLS as the mouse was running to inhibit the activity of direct pathway D1-SPNs during task learning. An example trace from a DMS implanted mouse shows significant reduction in calcium signals in D1-SPNs as the drug is injected, while the red wavelength rhodamine signal increased (Supplementary Figure 3), confirming local infusion.

All mice, independent of the region of injection, showed delayed improvement in performance until day 5. By day 6 performance was quickly improved (Figure 3A). The coordination scores for days 2 through 5 were significantly lower for animals injected with SCH39166 in the DMS (2-way ANOVA Time x Column Factor *p*<0.05, *Bonferroni’s multiple comparisons test* for day 02 \*\**p*<0.01, day 4 \**p*<0.05, day 5 \*\**p*<0.01), as compared to saline injected animals. Interestingly, injections of the D1-antagonist in the DLS also slowed learning, though it only reached significance for day 5 (2-way ANOVA Time x Column Factor *p*<0.05, *Bonferroni’s multiple comparisons test* for day 5 ^#^*p*<0.05).

**Figure 3.**
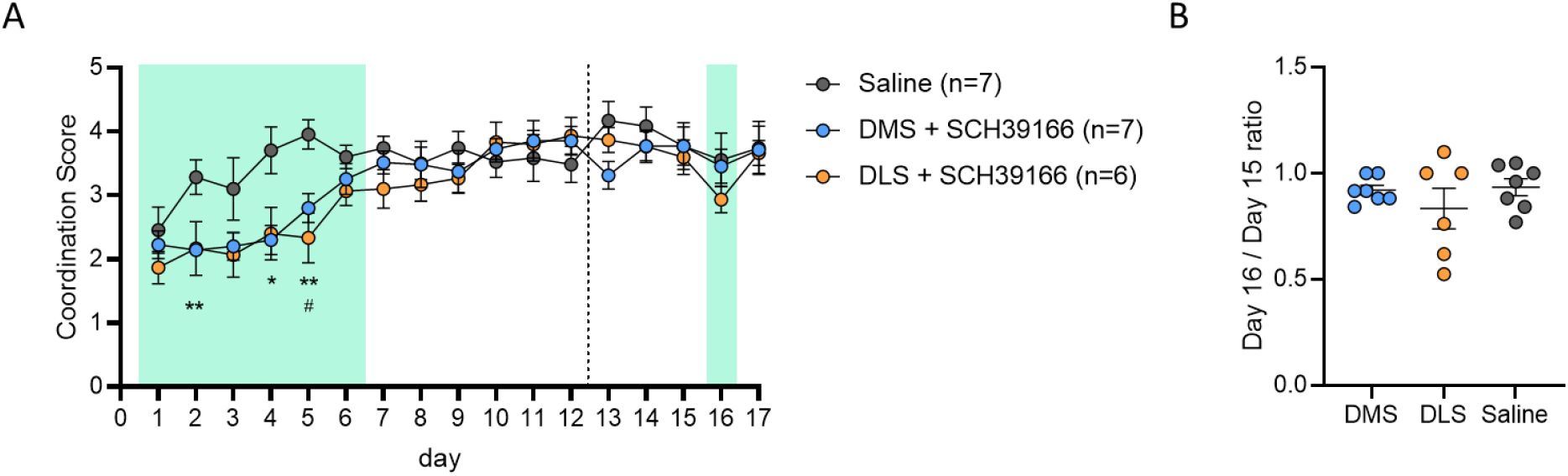
Local D1 neuron inactivation in DMS or DLS during the Treadmill Training Task. Mice were locally injected with a solution of SCH39166, a D1-antagonist. All animals were injected for the first 6 days of the task (injection running for the first 3 minutes of the test each day). No drug was injected from day 7 to day 12. Mice were then left to rest for 7 days and tested again for 5 consecutive days (day 13 to day 17). Another injection of SCH39166 was administered on day 16. Injection days are indicated by green shading. A) D1 antagonism in the DMS caused delayed improvement in the performance, as shown by low coordination score from day 2 to day 5 when compared to saline injected animals (2-way ANOVA Time x Column Factor *p*<0.05, *Bonferroni’s multiple comparisons test* for day 02 \*\**p*<0.01, day 4 *p*<0.05, day 5 \*\**p*<0.01). Similar to DMS-injected animals, application of the same D1 antagonist to the DLS altered performance, with a delayed improvement of the coordination score particularly on day 5 (2-way ANOVA Time x Column Factor *p*<0.05, *Bonferroni’s multiple comparisons test* for day 5 ^#^*p*<0.05). B) After a week without treadmill, both DLS and DMS-injected animals performed similarly to the previous running day. Conversely to DMS-injected mice, DLS-injected animals showed a decrease in performance after a single injection on day 16 (1-way ANOVA p<0.01, *Bonferroni’s multiple comparisons test* p=0.47), but quickly recovered the following day with no injection. Injection of SCH39166 in the DMS on day 16 did not affect performance as compared to control animals (1-way ANOVA p=0.85).

To investigate DMS and DLS contributions to performance once the skill is acquired, we recorded 5 additional days one week after the first session (labeled day 13 to day 17). On day 16, mice were injected with the same dose of SCH39166 as previously. Interestingly, injection in the DLS decreased the coordination score that however did not reach significance by *post-hoc* test (Figure 3A & B; 1-way ANOVA p<0.01, *Bonferroni’s multiple comparisons test* p=0.47). This decrease recovered the following day when mice were tested again without the D1-antagonist. Importantly, inhibition of DMS on day 16 showed no impairment in performance (Figure 3A & B; 1-way ANOVA p=0.85). No apparent changes in coordination score on day 16 for DMS injected mice suggest that this region is highly relevant in early training but is less necessary for performance of the previously learned tasks.

## Discussion

Current knowledge of motor learning comes from studies analyzing simple tasks, such as lever presses to achieve a food or drug reward or counting the number of falls from a motorized rotarod. While much has been learned from these approaches, they may not be applicable to learning more complex skills, such as changes in motor coordination during practice, a feature of activities ranging from athletic tasks to performance on a musical instrument. The training protocol introduced here, referred as the Treadmill Training Task, provides a means to activity of the direct pathway in the medial and lateral portions of the dorsal striatum while analyzing complex motor activity in freely moving animals in a carefully controlled locomotor task. To our knowledge, this is the first paradigm to measure changes in the activity of identified neurons during the learning of a complex and extended task requiring improvements in motor coordination. We furthermore developed a computational method to interpret multi-limb behavioral data and identify patterns of motor function that contribute to the improvement of coordination of a running mouse during practice. In doing so, we found that DMS and DLS activity progresses differently as animals gain skill in a motor task, revealing neuroanatomic specificity in progressive motor learning.

Our results are consistent with the hypothesis that DMS activity might decrease during complex motor learning (Cataldi et al., 2021). Similarly, a recent investigation shows that corticostriatal postsynaptic responses in the DMS undergo a transient decrease after training on the rotarod task, although those recordings were conducted in brain slice preparations and could not measure changes in activity during the task training (Badreddine et al., 2022).

Although we found that the amplitude of calcium events in the DMS changes over training, the rate of calcium activity transients did not. Our results are consistent with a report by Wightman et al. that demonstrated distinct active and silent regions in the striatum that exhibit different SPN activity and dopamine release patterns as a task is acquired (Owesson-White et al., 2016), a feature further consistent with the recent study on brain slices from trained animals (Badreddine et al., 2022). Badreddine’s study suggests that a reorganization of high activity cells increases the signal-to-noise ratio and contributes to better transmission of information. In this regard, we note that calcium signals detected by fiber photometry describe the bulk activity of a multitude of neurons, and an overall reduction in amplitude but not in the number of events could indicate that fewer, “specialized” neurons fire at the same rate once the skill is acquired (Bamford, Wightman, & Sulzer, 2018). Future experiments may explore this hypothesis by examining single cell activity over training using imaging techniques that utilize miniature head-mounted microscopes.

An important advantage to our approach is the ability to expose the regions being recorded to local pharmacological manipulations. Such experiments demonstrate that inhibition of D1 receptors delayed motor coordination learning in both the DMS and the DLS. The ability of mice to still learn improved performance on the Treadmill Training Task, albeit with a substantial lag compared to saline-treated mice, suggests that additional regions or cell types may promote motor learning with a lower efficiency in the absence of striatal activity. Interestingly, while no change in DLS activity was detected during training, inhibition of D1 dopamine receptors in the DLS later in training affected performance after the motor skills had been acquired. Similarly, Badreddine and colleagues did not find changes in mean amplitude nor overall changes in percentage of high activity cells in DLS brain slices due to training, although they detected a spatial reorganization of clusters of high activity cells in this region in late training (Badreddine et al., 2022). Such findings, combined with our pharmacology results, suggest that the DLS is particularly important for the expression of learned skills. Dysfunction of dopamine signaling to DLS SPNs may affect the neuronal organization required for proper performance of a learned skill, causing the decrease in coordination in the Treadmill Training Task we observe after local application of the D1 antagonist.

Conversely, inhibition of DMS D1-SPNs did not decrease coordination scores after learning, consistent with the idea that later in training the DMS is disengaged and no longer necessary for expression of the skill (Badreddine et al., 2022; Cataldi et al., 2021). In summary, these results suggest that dopaminergic regulation of activity of D1-SPNs in the DMS is required for proper skill acquisition, while DLS direct pathway neurons contribute to performance in both early and late training.

Given the complexity of the circuits involved in motor learning and the multiple behavioral variables of each task, caution is required when interpreting the animal behavior and the interaction between the different brain regions. Dopamine neurons display different patterns of activation in different behavioral tasks, e.g., in head fixed vs. freely moving animals or depending on whether a reward is present (Coddington & Dudman, 2019) and the activity of striatal neurons may be influenced by similar considerations (Cataldi et al., 2021). We note that the treadmill protocol could encompass aspects of aversion, as the mouse attempts to avoid hitting the back of the box, although we observed no change in calcium signals when the mouse was placed in the box, arguing against a classical fear response. A similar approach could be developed to examine spontaneous running on a wheel, which would avoid box confinement and the forced component of running given by the motorized system.

The Treadmill Training Task provides a means to evaluate motor ability in healthy animals and disease models and is designed to be useful for the evaluation of motor disorders and motor learning deficits due to neurodevelopmental disease, drug dependence, neurotoxic regimens, stroke, seizures, or injury. This work will assist in the development of more efficient therapies for these disorders.

## Acknowledgements

We sincerely thank the efforts of Dr. Moshe Shalev and the staff at the Division of Comparative Medicine, Columbia University, for advice and training in animal husbandry. We thank Vanessa Morales for breeding and care of all animals used in this study.

## Contributions

SC conceptualization, project design and administration, investigation, data curation, formal analysis, writing-original draft, editing

CL conceptualization, project design and administration, writing-original draft, editing

SN data processing and analysis, editing

GK data analysis, editing

DS conceptualization, provided direction for the experimental approach, resources, supervision, wrote and critically revised the manuscript, obtained funding.

---

**Supplementary Figure 1.**
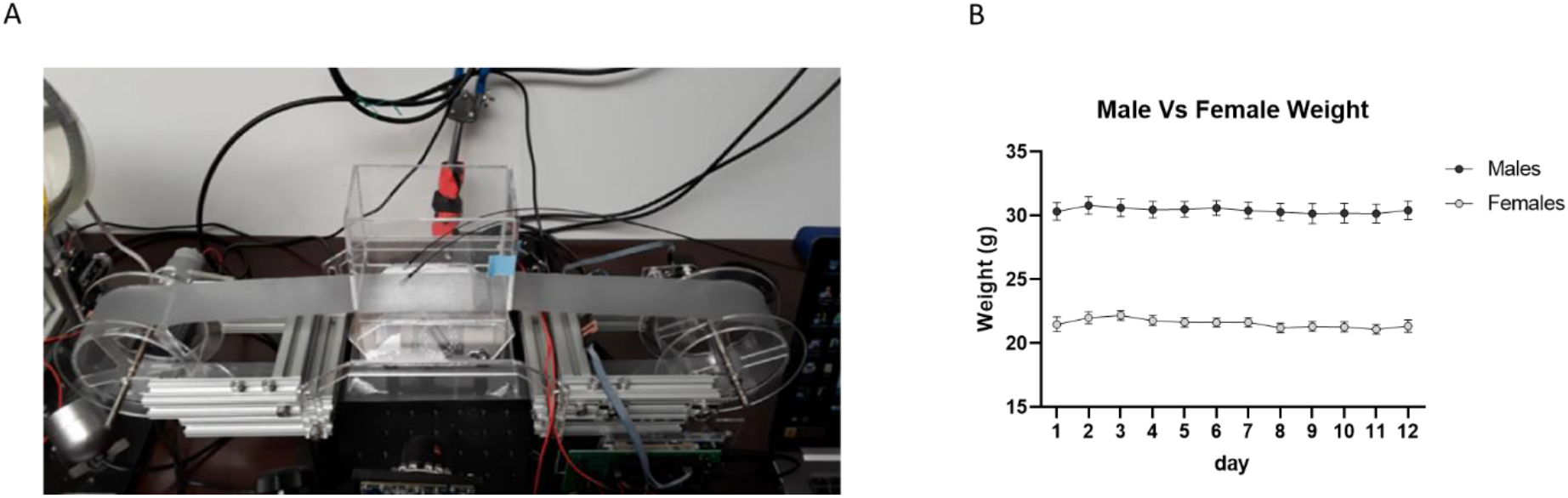
A) The custom-built treadmill system. A clear circular belt is moved by a motor controlled by a microcontroller. A clear acrylic box is placed around the running area to prevent the mouse from jumping off the treadmill during the experiment. A mirror is placed below the treadmill to visualize movement of all four paws during running via a camera located on the side (not in picture). B) Mouse weight was recorded before each day of experiment. 12 consecutive days of training did not affect mouse weight (1-way ANOVA *p*=0.90 for time). Data showing differences between male and female weight at this age (3-4 months of age; n=11 males; n=7 females; 2-way ANOVA *p*<0.0001).

**Supplementary Figure 2.**
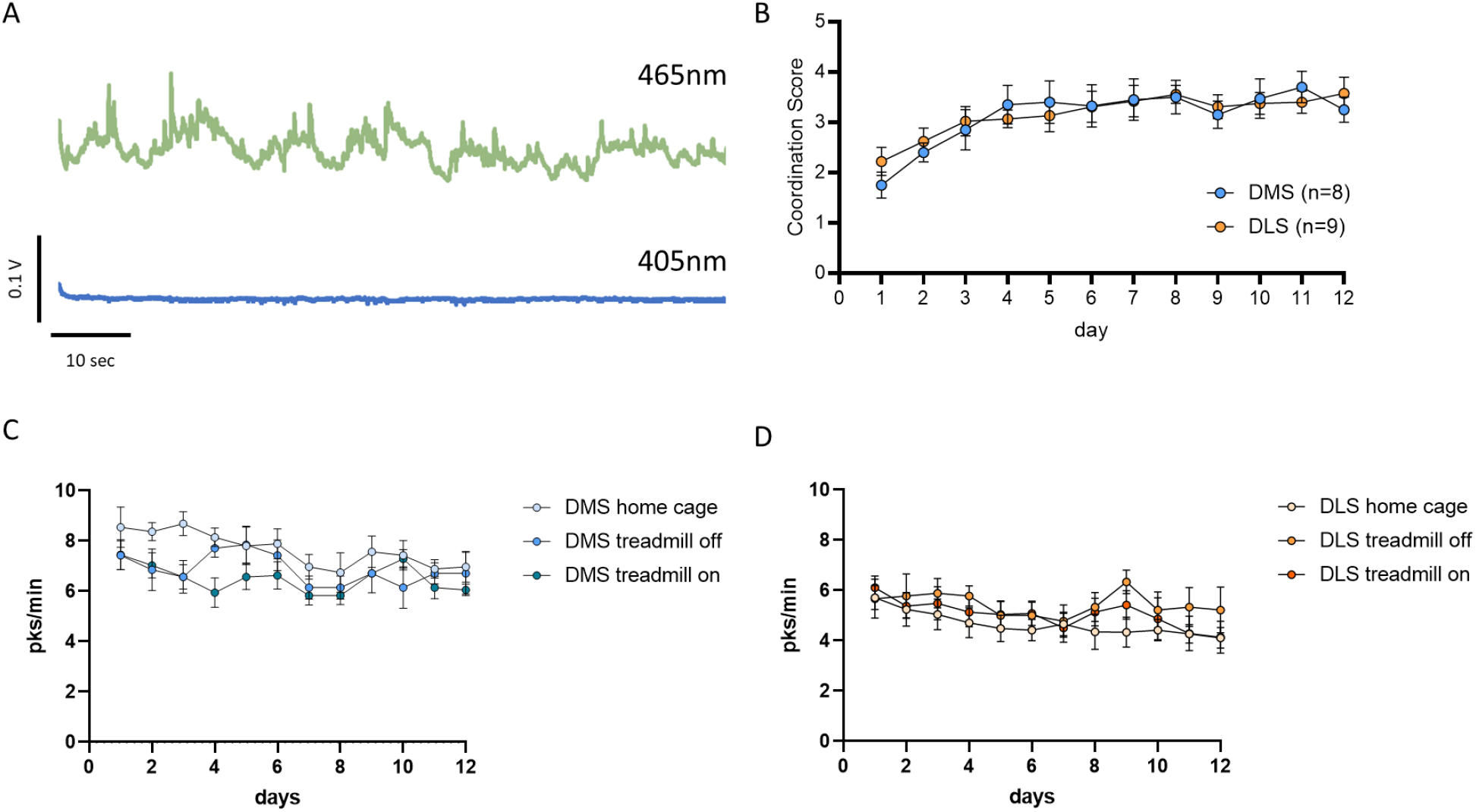
A) Raw sample fluorescence traces recorded during baseline with excitation at 405 nm (bottom trace), GCaMP6f isosbestic point, in comparison to calcium signal at 465 nm (top). B) Coordination score from treadmill running for DMS and DLS injected animals. Mice performed significantly better within 2-4 days from day 1 of training, indicating that surgery, viral expression, and optic fiber implant did not affect running abilities and motor learning (1-way ANOVA \**p*<0.05 for DMS mice and \*\**p*<0.01 for DLS mice). C,D) Average calcium event rate for D1-SPNs in the DMS (C) and DLS (D). Measurements for epochs of running time (treadmill on), time in which the treadmill is off, and event counts in home cage baseline recording were overlapped. There were no significant changes in the average event rate over the 12 days of training for DMS (2-way ANOVA Time x Column Factor *p*=0.82) or DLS recording (2-way ANOVA Time x Column Factor *p*=0.77 for DLS).

**Supplementary Figure 3.**
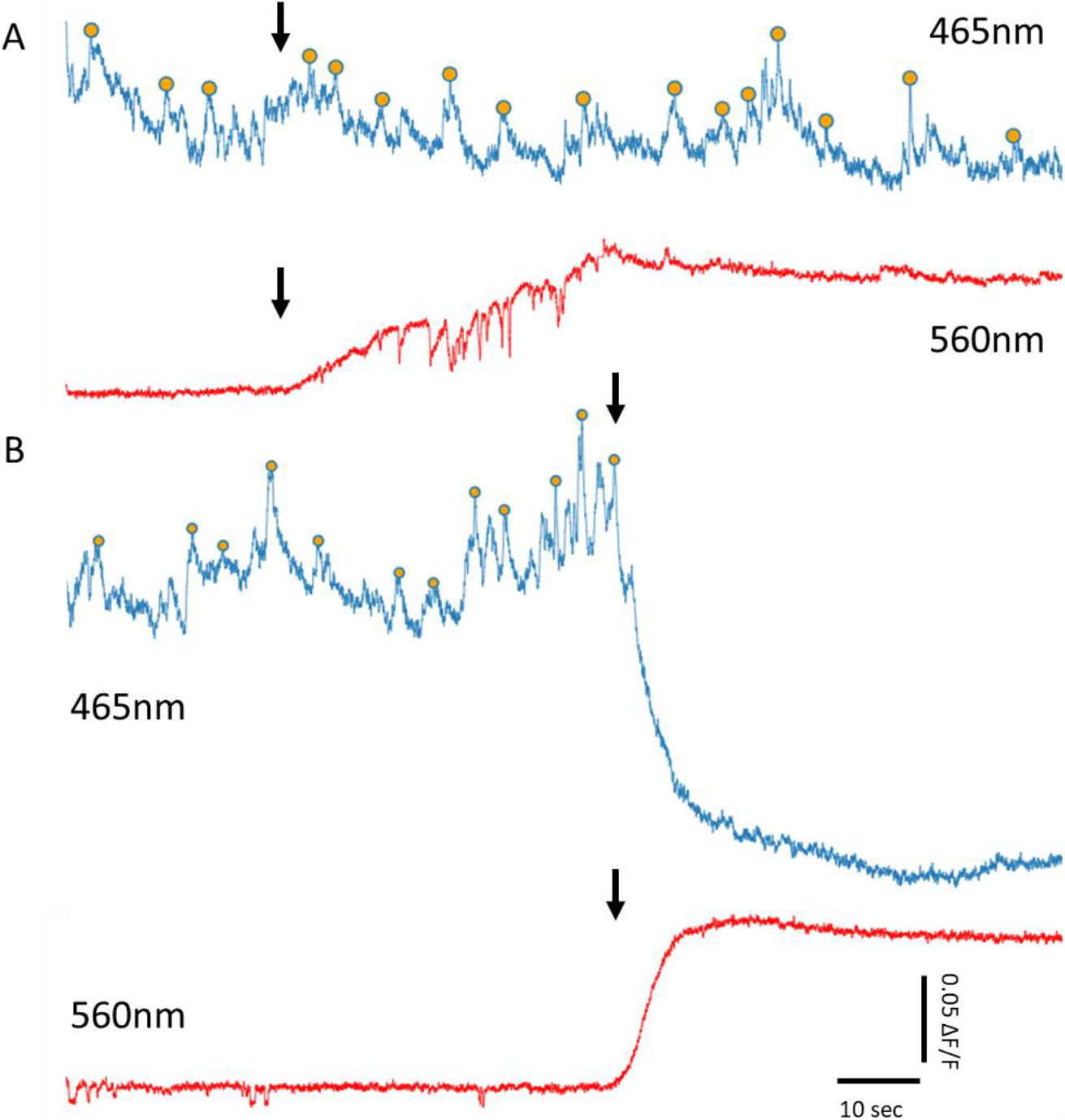
A) Representative calcium trace (in blue, detected events indicated by orange dots) of D1-SPNs GCaMP6f signal in the DMS of a freely moving mouse, while injected with a control solution of saline + rhodamine. Rhodamine signal is recorded at 560 nm to confirm injection local to the photometry fiber (red). Local injection did not alter calcium signals in this control. Injection time indicated by arrows. B) Similar recording of D1-SPNs in the DMS during injection of a D1-antagonist (SCH39166 0.2mg/Kg), in solution with rhodamine. Injection of the antagonist significantly reduced the baseline calcium signal and eliminated calcium events in the DMS.

**Supplementary Figure 4.**
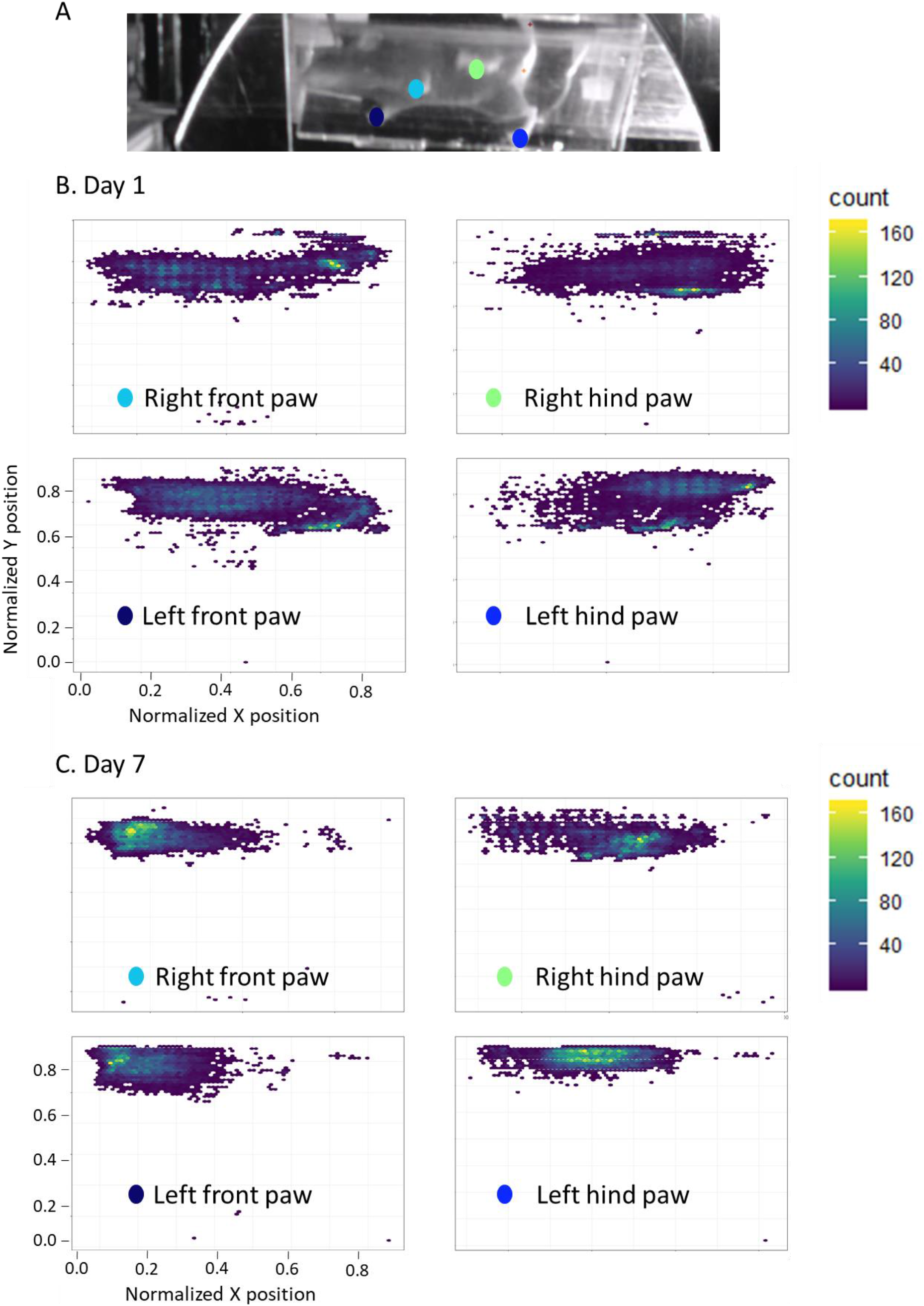
Heatmaps of paw placement data obtained from DeepLabCut for one mouse on day 1 (B) and day 7 (C) of the Treadmill Training Task. Each heatmap represents data from a different paw and is color-labeled as per the schematic in A and in Figure 1B (top left for the right front paw – light blue dot in panel A; bottom left for the left front paw – dark blue dot in panel A; top right for the right hind paw – green dot in panel A; bottom right for the left hind paw – blue dot for panel A). X and Y axis values obtained from tracking are normalized. Each heat map shows likelihood of placing a specific paw in a specific set of coordinates. The blue color represents regions with low number of paw placements, while yellow indicates high chance of placing in those coordinates. B) Heatmaps for one control mouse on day 1 of training. The spread of paw placement indicates poor ability of the mouse to run into place. While running the mouse was placing its paws all over the 2D field and rarely in the same spot. C) Heatmaps for the same control animal on day 7 of training. This data show more consistent paw placement with the mouse stepping mostly in the same area. This shows the ability of the mouse to run in place, indicating improved proficiency in running.

**Supplementary Figure 5.**
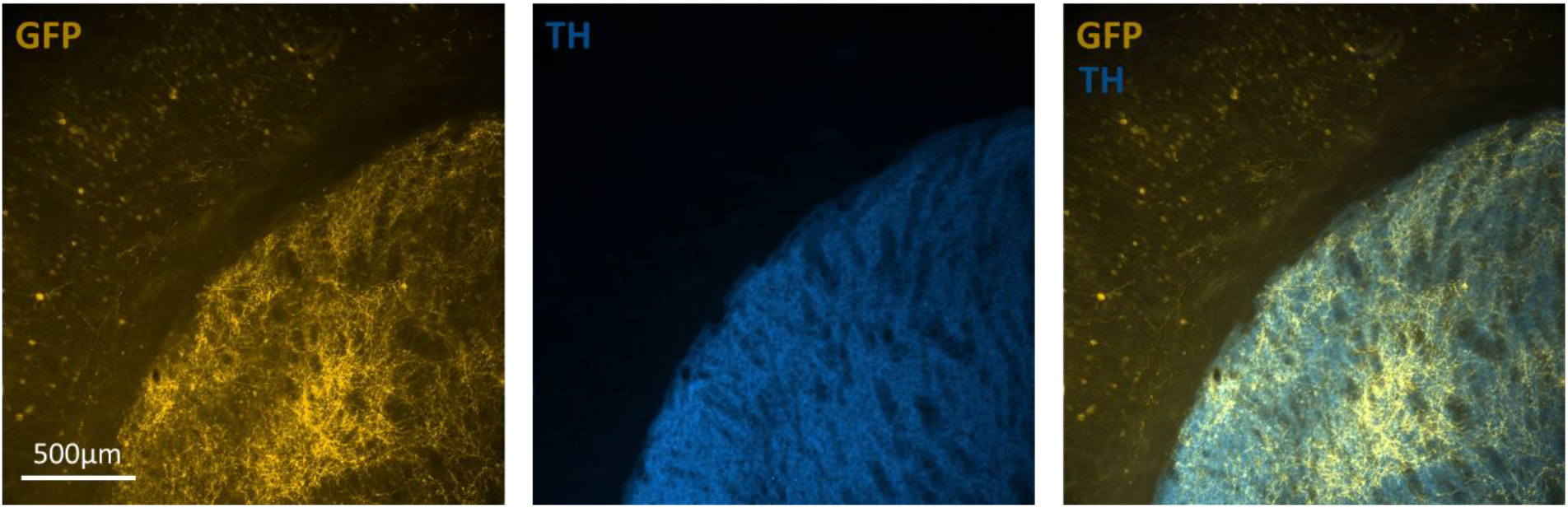
Sample micrographs of coronal sections of the dorsal striatum of D1-cre animals injected with AAV vectors containing Flex-GCaMP6f. GCaMP6f expression is confirmed by GFP immunolabeling (yellow) co-stained with TH antibody (blue) to visualize the striatum.

